# Radular teeth matrix protein 1 directs iron oxide deposition in chiton teeth

**DOI:** 10.1101/2024.11.02.621658

**Authors:** Michiko Nemoto, Koki Okada, Haruka Akamine, Yuki Odagaki, Yuka Narahara, Kenji Okoshi, Kiori Obuse, David Kisailus, Hisao Moriya, Akira Satoh

## Abstract

Chitons deposit magnetite, a type of iron oxide, on the cusps of their radular teeth. The molecules that control the deposition of iron oxide on chiton teeth have not been identified. Using immunofluorescence staining, we found that chiton-specific radular teeth matrix protein 1 (RTMP1) is highly expressed in tooth epithelial cells prior to iron oxide deposition and preexists in the cusp at the site of iron oxide deposition. RTMP1 was found to be secreted from the elongated epithelial cells covering teeth and transported into teeth through microvilli. Moreover, recombinant RTMP1 exhibited the ability to form iron oxide. These results indicate that RTMP1 is a key molecule for iron oxide deposition in chiton teeth.

## Introduction

Living organisms can synthesize biominerals with a variety of remarkable properties under physiological conditions. These highly functional biominerals are synthesized through precise formation processes controlled by biomolecules (*1–7*). Chitons, mollusks found in rocky coastal areas, form magnetite (Fe_3_O_4_) teeth that exhibit greater wear resistance than zirconia, making them an excellent model for studying such biomineralization (*8–10*). Chitons have a feeding organ called a radula and deposit magnetite on the cusps of their radular teeth (Fig. 1A). Chiton teeth were originally discovered by Lowenstam in 1962 as the first biologically formed magnetite (*11*). Chitons use these tough teeth to scrape algae and other organisms from rocks for food. As the magnetite-deposited cusps wear and fall off during feeding, they are replaced by newly formed teeth pushed forward from behind. Therefore, new teeth are continuously formed in the radular tissue, and teeth at different maturation stages are present in one radula (Fig. 2A) (*12*). The formation of chiton teeth is regulated by the epithelial cells covering the teeth in a tissue called the radular sac. Tooth formation begins with the formation of a transparent tooth matrix composed of α-chitin and protein by odontoblast cells at the posterior end of the sac. Subsequently, mineral aggregates of amorphous iron oxide (ferrihydrite) are deposited on the chitin fibers, resulting in reddish-brown teeth. The amorphous iron oxide then crystallizes to form magnetite, resulting in black teeth (Fig. 2A). Previous studies have indicated the existence of a protein that causes iron oxide deposition in the cusp of chitons, but this protein remains unidentified (*13, 14*).

**Fig. 1.**
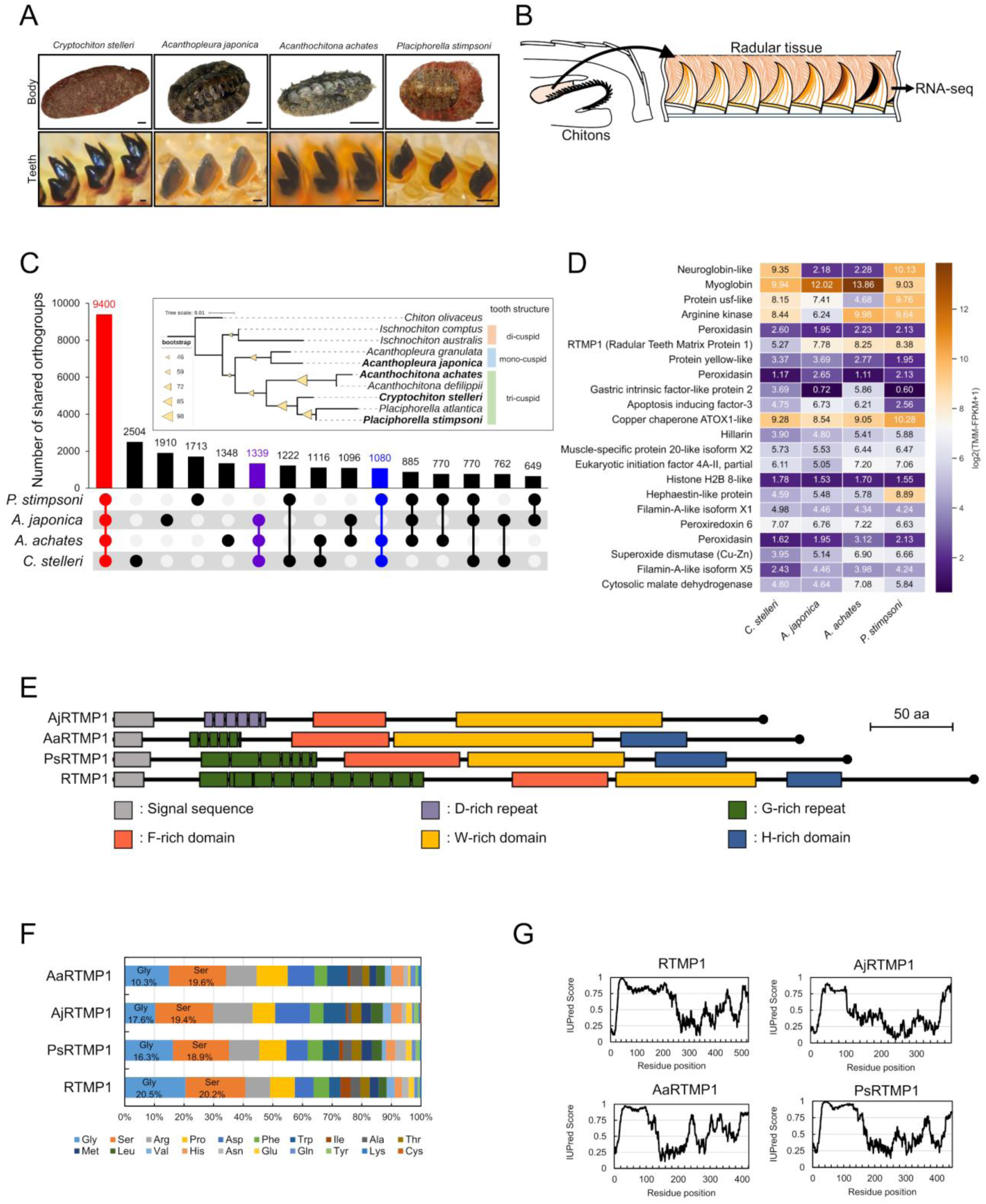
A transcriptomic comparison of radular tissues revealed that RTMP1 and its homologs are chiton-specific proteins. (**A**) Chitons used in this study. The upper images show the whole body, and the lower images show the teeth. Scale bars, 1 cm (top images) and 100 μm (bottom images). (**B**) Schematic of the RNA-seq experiments. (**C**) UpSet plot showing shared and unique orthologous protein groups (orthogroups) across four chiton species. The numbers at the top of the bars indicate the number of orthogroups shared among different chiton species. The shared orthogroups among all the chitons analyzed in this study are highlighted in red, those among herbivorous chitons are highlighted in purple, and those among chitons with tricuspid teeth are highlighted in blue. The inset shows a phylogenetic tree of CO1 sequences of chitons. The species used in this study are shown in bold. Information on the tooth shape is shown on the right. (**D**) Heatmap showing the gene expression profiles of homologs of mineralized cusp-specific proteins, originally identified in *C. stelleri*, across four chiton species. (**E**) Schematic of the primary structures of RTMP1 and its homologs. (**F**) Comparison of the amino acid composition of RTMP1 and its homologs. (**G**) Disordered region prediction for RTMP1 and its homologs using IUPred.

**Fig. 2.**
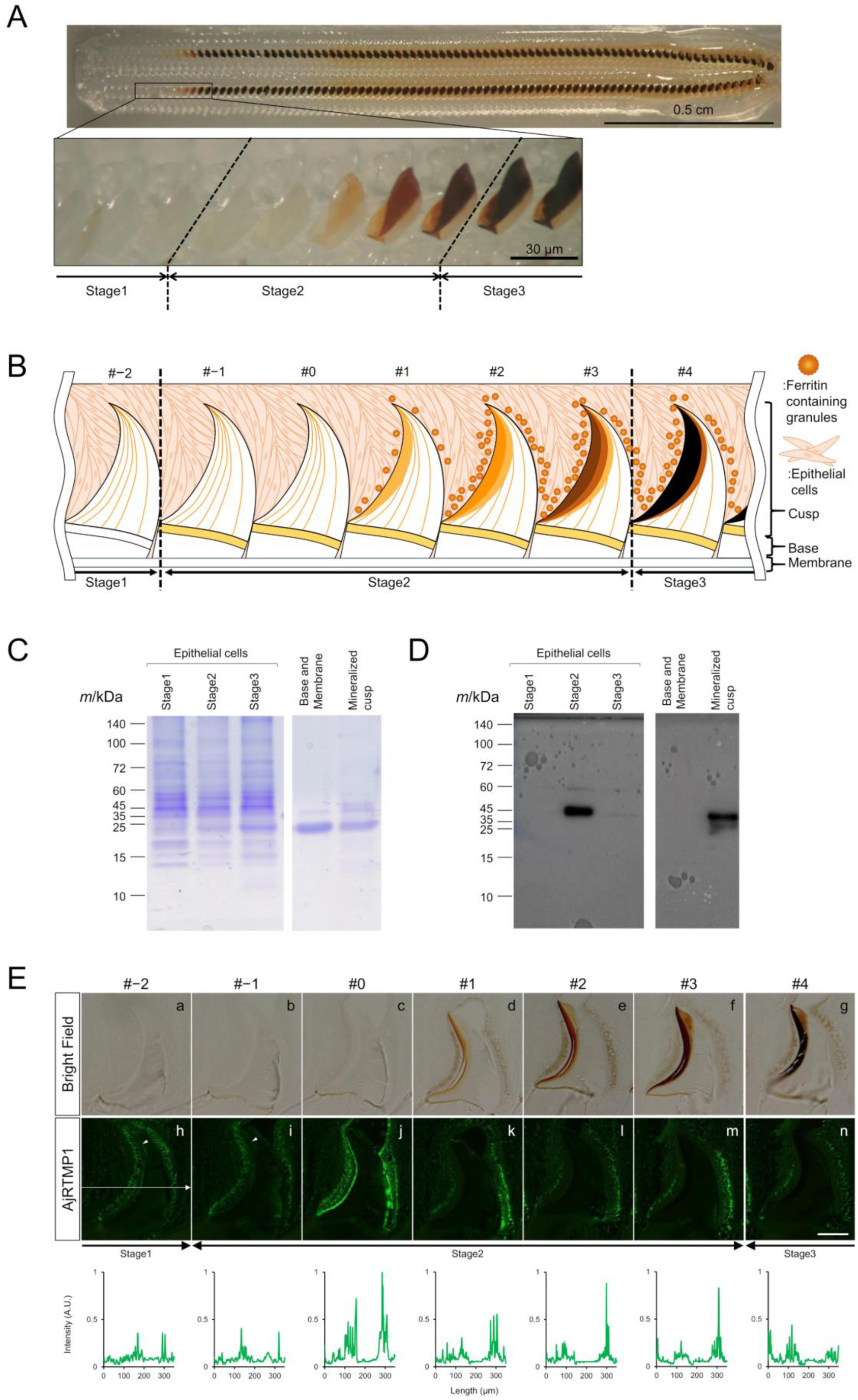
Localization of AjRTMP1 during tooth maturation. (**A**) Radula of *A. japonica* with teeth in different stages of maturation. (**B**) Schematic representation of the radular tissue. (**C**) SDS‒PAGE and (**D**) Western blot of proteins extracted from *A. japonica* radular tissue. Approximately 10 μg of protein extracted from epithelial cells at stage 1, stage 2, and stage 3, as well as proteins extracted from the base, membrane, and mineralized cusp after removal of epithelial cells, was subjected to tricine SDS‒PAGE and stained with Coomassie blue. Proteins fractionated by tricine SDS‒PAGE were transferred to a polyvinylidene fluoride (PVDF) membrane, followed by immunoblotting with polyclonal anti-AjRTMP1 antibodies. (**E**) The 4 µm paraffin sections of *A. japonica* radular tissues were stained with an anti-AjRTMP1 antibody followed by Alexa Fluor 488-conjugated anti-rabbit IgG. The tooth with the earliest visible mineral deposition on the cusp was designated tooth #1. Teeth with up to three immature rows and up to three rows more mature than tooth #1 were analyzed. Brightfield (a-g) and fluorescence (h-n) images of (a, h) tooth #−2, (b, i) tooth #−1, (c, j) tooth #0, (d, k) tooth #1, (e, l) tooth #2, (f, m) tooth #3 and (g, n) tooth #4. The graphs in the bottom panel are the fluorescence profiles of AjRTMP1 (green) along the direction of the arrows in the fluorescence image (h). For comparison of the changes in fluorescence intensity between the images, normalization was performed by dividing by the maximum brightness value across all images. The white arrowheads highlight the localization of AjRTMP1 within the teeth. Scale bar, 100 μm.

To elucidate the mechanism of tooth biomineralization, we previously performed a transcriptomic analysis of the radular tissue of the gumboot chiton, *Cryptochiton stelleri*, and identified genes expressed in the radular tissue (*15*). In addition, proteins extracted from mineralized cusps were compared with those extracted from base and radular membranes, and 22 proteins specific to mineralized cusps were identified (*15, 16*). These proteins are believed to be important for iron oxide deposition on teeth, but their functions remain unclear.

## Results

### RTMP1 and its homologs are chiton-specific proteins

To identify proteins related to magnetite mineralization present across chiton species, we performed de novo assembly of transcriptomes of radular tissue from three chiton species, namely, *Acanthopleura japonica*, *Acanthochitona achates* and *Placiphorella stimpsoni,* and performed cross-species comparisons with the previously analyzed transcriptomes of *C. stelleri* (Fig. 1AB, table S1). A total of 9,400 orthologous protein groups were found to be shared among the four chitons (Fig. 1C). Among them, homologs of 22 previously identified mineralized cusp-specific proteins in *C. stelleri* were found from the transcriptomes of the other three chiton species (Fig. 1D, table S2). Among the 22 mineralized cusp-specific proteins, RTMP1 shared no significant homology with proteins from other organisms and was considered a chiton-specific protein. Additionally, a homolog of RTMP1 was found in the genome of the chiton *Acanthopleura granulata* (*17*). The RTMP1 homologs in *A. japonica*, *A. achates*, and *P. stimpsoni* were named AjRTMP1, AaRTMP1, and PsRTMP1, respectively (Fig. 1E). All of these sequences contain the predicted signal peptides and a high percentage of serine residues (˃19%) (Fig. 1F). Sequence alignment revealed that the Gly-, Phe-, Trp- and His-rich domains are conserved among RTMP1, AaRTMP1 and PsRTMP1 (Fig. 1E, fig. S1). AjRTMP1 does not contain Gly- or His-rich domains, but it does contain an Asp-rich domain. Among the previously predicted chitin-binding residues in RTMP1, only Trp is conserved in AjRTMP1, AaRTMP1, and PsRTMP1 (fig. S1). A single Trp conserved in four RTMP1s has been reported to play a key role in chitin binding in *Bacillus circulans* chitinase A1 (*18*). Sequence alignment revealed that the sequence similarity among RTMP1, AaRTMP1 and PsRTMP1 is 51.7–75.3%. Whereas, AjRTMP1 presents low sequence similarity (maximum 35.8%) with the other three RTMP1 homologs (fig. S1). On the basis of structural prediction from the amino acid sequence, the first half of all RTMP1 homologs have a region predicted to be disordered (Fig. 1G). Mass spectrometric analysis of proteins extracted from *C. stelleri* radular tissue confirmed that a serine residue in the SSSE motif of RTMP1 is phosphorylated (fig. S2). Notably, the SSSE motif of RTMP1, which was found to be phosphorylated, is present in all the RTMP1 homologs of the chiton species (fig. S1).

### AjRTMP1 is expressed in the radular tissue at the onset of iron deposition and eventually localizes to the tooth cusps

Next, we examined AjRTMP1 protein expression in *A. japonica* radular tissue using an anti-AjRTMP1 antibody. In this analysis, the radular tissue was divided into three stages: stage 1 (8–9 teeth), consisting of transparent teeth composed mainly of chitin; stage 2 (3–5 teeth), consisting of teeth with a reddish-brown color due to iron deposition; and stage 3 (10 teeth), where amorphous iron oxide crystallized to form magnetite, resulting in black teeth (Fig. 2A and B). In stage 2, iron is deposited first at the junction zone of the cusps and bases and then at the leading edge of the cusps. We designated the first tooth where iron deposition occurred at the leading edge as #1; designated less mature teeth as #0, #−1, and #−2; and designated more mature teeth as #2, #3, and #4 (Fig. 2B). Proteins were extracted from epithelial cells at each stage, and Western blotting was performed. As a result, a band at approximately 43 kDa, which is the protein size expected from the amino acid sequence of AjRTMP1, was detected in stage 2 epithelial cells (Fig. 2C and D). The results revealed that AjRTMP1 was most highly expressed in stage 2. This finding is consistent with the results of the qPCR analysis of *C. stelleri*, which revealed that the RTMP1 gene had the highest expression in stage 2 (fig. S3). Next, Western blotting was performed using proteins extracted from the mineralized cusps and the base plus radular membranes that support the cusps (*16*). As a result, a band corresponding to AjRTMP1 was observed in the cusp fraction (Fig. 2C and D). The band size for AjRTMP1 observed in the cusp fraction was found to be slightly smaller than the predicted size on the basis of its amino acid sequence (Fig. 2C and D). These results suggest that AjRTMP1 may undergo post-translational modifications after being secreted from epithelial cells before being localized to cusps. These results indicate that AjRTMP1 is also a mineralized cusp-specific protein, similar to RTMP1, which was originally identified from *C. stelleri* radulae as a mineralized cusp-specific protein (*15*). Taken together, these results indicate that RTMP1 and its homologs are expressed in stage 2 epithelial cells at the onset of iron deposition and eventually localize to the cusps.

### The timing and localization of AjRTMP1 expression are related to the timing and site of iron oxide deposition in the tooth cusps

To elucidate the localization of AjRTMP1 in radular tissues, we performed immunofluorescence staining using an anti-AjRTMP1 antibody. Sections of *A. japonica* radular tissues were prepared and stained with an anti-AjRTMP1 antibody and Alexa Fluor 488-conjugated anti-rabbit IgG. The results revealed that AjRTMP1 was initially expressed around teeth #−2 to #−1 and was expressed at the highest level around tooth #0 (Fig. 2E). Furthermore, AjRTMP1 was confirmed to be localized at the leading edge of the tooth (Fig. 2E, white arrowhead). In the epithelial cells of teeth #−2 to #0, AjRTMP1 was uniformly localized on both the leading edge and trailing edge of the tooth. In contrast, in the teeth following tooth #1, where iron oxide was deposited on the leading edge, AjRTMP1 was localized mainly to the epithelial cells of the trailing edge. These findings suggest that AjRTMP1 infiltrates from the trailing edge into the interior of teeth after iron deposition on the leading edge. Brown ferritin granules were observed around tooth #1 and the following teeth (Fig. 2E). Iron ions transported by ferritin may be deposited in the cusp because the timing of ferritin appearance in epithelial cells coincides with the timing of iron deposition in the tooth cusp (*19, 20*). On the basis of the above results, we propose the following hypothesis. In teeth #−2 to #0, the organic template AjRTMP1 is transported and localized within the cusp. In tooth #1 and subsequent teeth, iron ions released from ferritin likely interact with AjRTMP1, resulting in the deposition of iron oxide on the cusp.

### AjRTMP1 is delivered from epithelial cells to the interior of cusps through microvilli

For a more detailed analysis of AjRTMP1 localization in epithelial cells, we performed multiplex staining of longitudinal and transverse tissue sections stained with an anti-AjRTMP1 antibody (Fig. 3A). Alexa Fluor 594-conjugated phalloidin and Hoechst were used to stain F-actin and nuclei, respectively. AjRTMP1 was confirmed to be expressed in the columnar epithelial cells covering the teeth (Fig. 3B). The high density of actin near the cusps was suggested to be due to the microvilli covering the cusps (*19*). Furthermore, the fluorescence signal indicating the localization of AjRTMP1 was stronger in the epithelial cells than in the cusps (Fig. 3B). To further elucidate the localization of AjRTMP1 in detail, we performed an analysis of sections with immunohistochemical staining using confocal laser scanning microscopy (Fig. 3C and D). The results confirmed that AjRTMP1 is localized in a linear pattern adjacent to the cusp at both the tip and base of the F-actin filaments. We also confirmed that AjRTMP1 is localized between the F-actin filaments (Fig. 3C). Immunostaining and confocal microscopy of transverse sections of radular tissue revealed that AjRTMP1 is localized within epithelial cells and at the leading edge inside the tooth cusps (Fig. 3D). Furthermore, prior to iron deposition in the cusps, AjRTMP1 was observed to enter the interior of the tooth from both the leading and trailing edges (Fig. 3D). However, after iron deposition in the cusps, AjRTMP1 was confirmed to flow mainly from the trailing edge (Fig. 3D). This finding is consistent with the results from the immunohistochemical analysis of the longitudinal section in Fig. 2E. Furthermore, after iron deposition occurred at the leading edge of the cusp, the localization of AjRTMP1 shifted slightly toward the trailing edge (Fig. 3D). These findings suggest that after iron oxide is formed at the leading edge with AjRTMP1 as a template, AjRTMP1 flows again into the immediate posterior region of the formed iron oxide, where new iron oxide is formed.

**Fig. 3.**
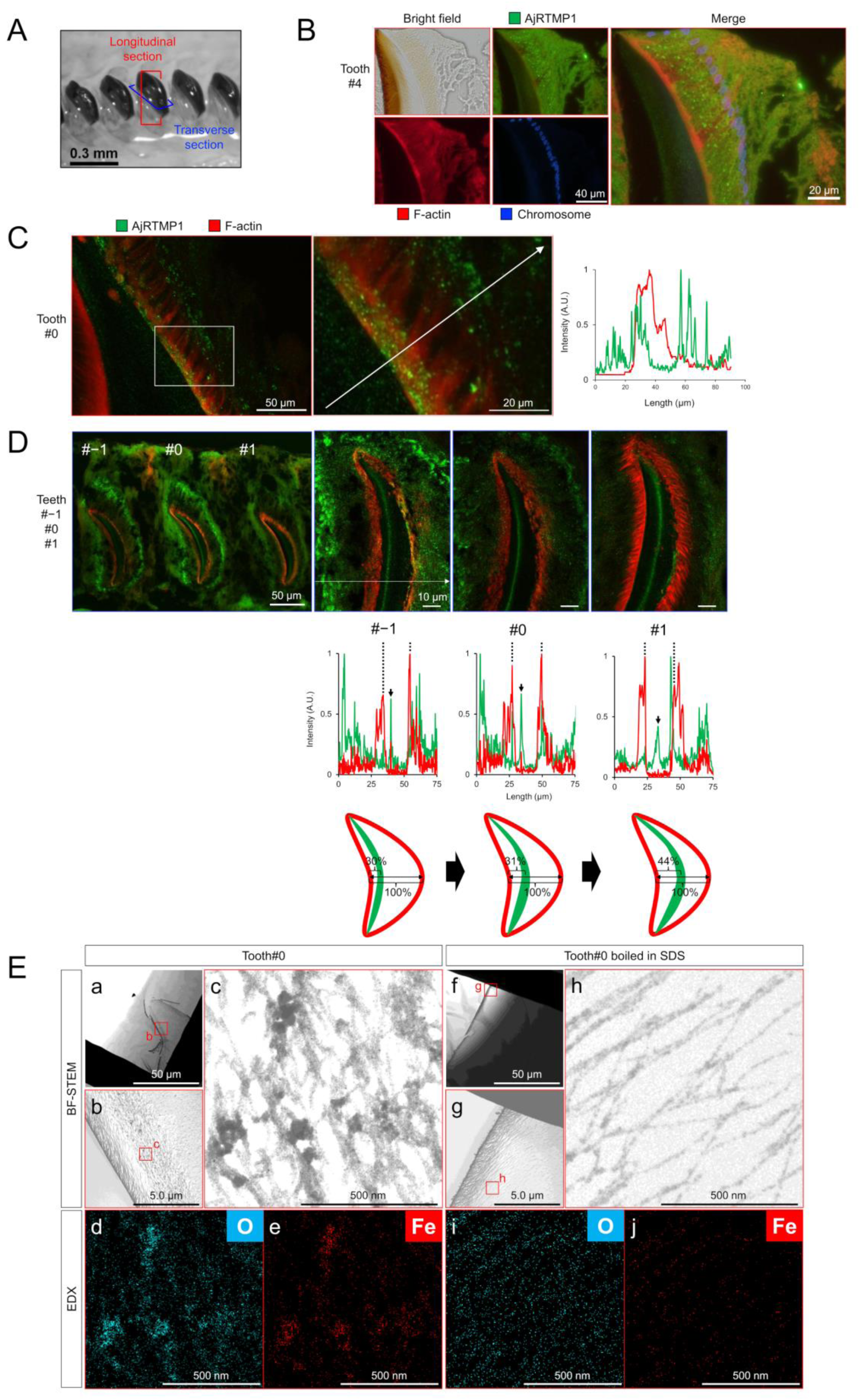
Multiplex immunofluorescence staining was performed on sections of *A. japonica* radular tissue. (**A**) Longitudinal and transverse sections of *A. japonica* radular tissue were prepared and used for analysis. (**B**) The 4 µm paraffin sections of *A. japonica* radular tooth #4 were stained with anti-AjRTMP1 antibody followed by Alexa Fluor 488-conjugated anti-rabbit IgG. Chromosomes (blue) and F-actin (red) were counterstained with Hoechst 33342 and Alexa Fluor 594 phalloidin, respectively. (**C**) A 20 µm longitudinal cryosection of *A. japonica* radular tooth #0 was stained with anti-AjRTMP1 (green) and Alexa Fluor 594 phalloidin (red) and analyzed using confocal microscopy. The graphs in the right panel are the fluorescence profiles of AjRTMP1 (green) and F-actin (red) along the direction of the arrows in the fluorescence image. (**D**) A 20 µm transverse cryosection of *A. japonica* radular teeth #−1, #0, and #1 was stained with anti-AjRTMP1 (green) and Alexa Fluor 594 phalloidin (red) and analyzed using confocal microscopy. The graphs in the bottom panel are the fluorescence profiles of AjRTMP1 (green) and F-actin (red) along the direction of the arrows in the fluorescence image. The lower schematic shows that as the tooth matures and mineralization occurs at the leading edge, the localization of AjRTMP1 shifts to the trailing edge. The peak indicated by the black dotted line was defined as the tooth surface, and the peak indicated by the black arrow was defined as the localization site of AjRTMP1. The relative position of AjRTMP1 within the tooth was then calculated. (**E**) In vitro mineralization experiments using radular teeth extracted from *A. japonica*. After removal of the epithelial cells, tooth #0 samples (a, b, c, d, e) and tooth #0 samples boiled in SDS (f, g, h, i, j) were both left in FeCl_3_ solution for 7 days and subsequently analyzed using TEM and EDX.

### RTMP1 binds to ferric ions and directs the formation of iron oxide

The immunohistochemical staining results suggested that AjRTMP1 directs the formation of iron oxide on chitin fibers as an organic template. Therefore, iron oxide formation experiments were performed using tooth #0, which presented the highest expression of AjRTMP1. After fresh radulae were extracted from the dissected specimens and epithelial cells were removed, the teeth were incubated in FeCl_3_ solution. As a result, iron oxide nanoparticle deposition was confirmed on the chitin fibers of the leading edge where AjRTMP1 was localized. After protein removal by SDS treatment, no particle deposition similar to that observed above was detected on the chitin fibers of the teeth (Fig. 3E). These results suggest that organic templates exist on the chitin fibers of the cusp prior to iron influx and that the iron released from ferritin infiltrates them, resulting in the deposition of iron oxide.

To determine whether RTMP1 acts as an organic template for iron oxide deposition, we used *Saccharomyces cerevisiae* to recombinantly express RTMP1 and purified the protein. RTMP1 without a signal sequence, which we named RTMP1 Δss, and RTMP1 without a G-rich repeat region, which we named RTMP1 ΔG, were expressed and purified as GST fusion proteins (Fig. 4A, fig. S4). The results of chitin-binding assays using colloidal chitin revealed that the recombinant GST-RTMP1 ΔG protein was found in the chitin-binding fraction, similar to the chitin-binding protein wheat germ agglutinin (WGA) (Fig. 4B). These results showed that the C-terminal region of RTMP1 has chitin-binding activity. In addition, the results of the iron binding assay using potassium ferrocyanide indicated that GST-RTMP1 Δss binds to Fe^3+^. The GST-RTMP1 ΔG protein showed attenuated iron-binding activity (Fig. 4C). Next, to evaluate whether RTMP1 has iron oxide crystal formation activity, we incubated recombinant GST- RTMP1 Δss with chitin nanofibers and FeCl_3_ solution (Fig. 4D). Scanning transmission electron microscopy (STEM) confirmed the formation of 27–53 nm particles on the chitin fibers. The results of STEM-energy-dispersive X-ray (EDX) mapping revealed the presence of iron and oxygen in the particle region, suggesting the formation of iron oxide particles on the chitin fibers. In the negative control experiment performed with GST-GFP, no iron oxide particles formed on the chitin fibers (Fig. 4E).

**Fig. 4.**
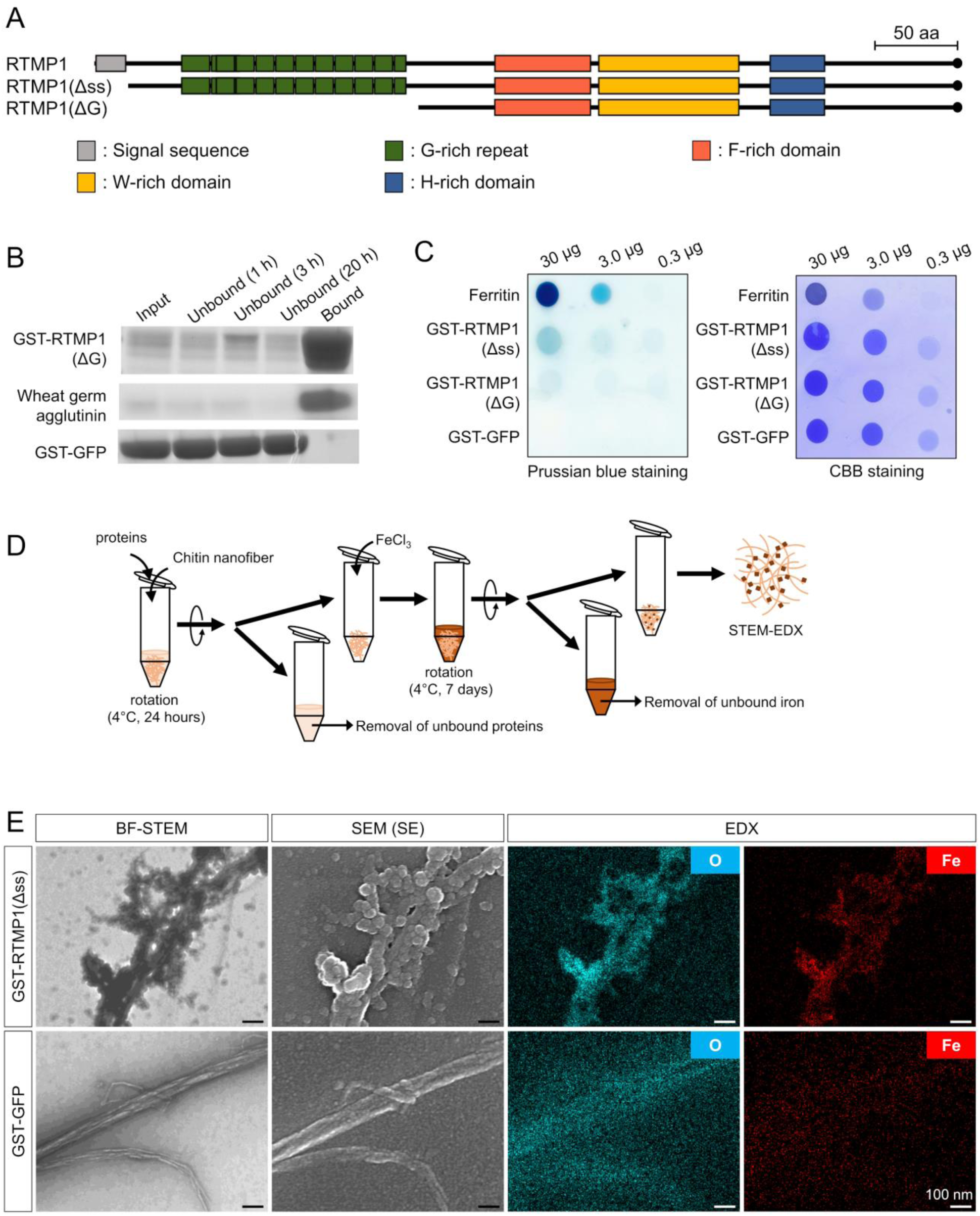
The recombinant RTMP1 exhibited iron ion binding and iron oxide precipitation activity. (**A**) Schematic primary structures of RTMP1 and recombinantly expressed RTMP1 Δss and RTMP1 ΔG. (**B**) Chitin binding assay using recombinant GST-RTMP1 ΔG. Recombinant GST-RTMP1 ΔG was incubated with insoluble colloidal chitin. Wheat germ agglutinin and GST-GFP were used as controls. After incubation for 1 h, 3 h and 20 h, the supernatants (unbound fractions) obtained together with the pellets (bound fractions) obtained after 20 h of incubation were collected and analyzed by SDS‒PAGE. (**C**) Iron binding analysis of recombinant GST-RTMP1 Δss and GST-RTMP1 ΔG. Prussian blue (left) was used to detect Fe(III), and CBB (right) was used to stain the proteins. Equal amounts of ferritin and GST-GFP were used as controls. (**D**) Schematic diagram of the iron oxide formation experiment. (**E**) STEM bright field images, SEM secondary electron images, and EDS maps (Fe and O) of iron oxide nanoparticles formed on chitin nanofibers in the presence of GST-RTMP1 Δss. GST-GFP was used as a control. Scale bar, 100 nm.

To investigate the function of RTMP1 in vivo, we performed gene knockdown experiments using RNA interference in *A. japonica* (fig. S5A). Two days after the injection of dsRNA for the AjRTMP1 gene into the body cavity, the mRNA expression level of AjRTMP1 in the radular tissue was examined.

Compared with the individuals injected with PBS or dsRNA for the GFP gene, the individuals injected with dsRNA for the AjRTMP1 gene presented ∼69% lower mRNA expression levels (fig. S5B). Microscopic examination of the radular tissue sections revealed that in the individuals injected with dsRNA for GFP, similar to WT individuals, iron was first deposited in tooth #1, followed by a clear progression of mineralization as maturation progressed, with the color gradually changing from light brown to dark brown and then to black. In contrast, in the individuals with decreased AjRTMP1 mRNA expression, after iron was deposited in tooth #1, teeth #2, #3, and #4 did not show the same clear progression of mineralization as described above, and the change in saturation values indicating color change was minimal (fig. S5C and D). These findings suggested that the reduction in AjRTMP1 expression prevented the progression of iron oxide deposition, resulting in the failure of gradual tooth maturation.

## Discussion

The iron-binding activity of GST-RTMP1 ΔG was lower than that of GST-RTMP1 Δss, suggesting the importance of the G-rich repeat region in iron-binding activity (Fig. 4C). Acidic amino acids have been reported to be critical for the function of the magnetite morphology-regulating protein Mms6 in magnetotactic bacteria (*21*). The N-terminal region of the G-rich repeat is rich in acidic amino acids, suggesting the potential importance of this region in interactions with iron ions (fig. S1). In addition, we observed phosphorylation of the G-rich region of recombinant GST-RTMP1 Δss (table S3), suggesting a potentially critical role of phosphate groups in the iron binding of this protein. Indeed, native RTMP1 was also phosphorylated (fig. S2). The importance of phosphate groups in binding to iron ions has been reported for other proteins (*22*). A previous amino acid analysis of the radular teeth of the chiton *Acanthopleura hirtosa* revealed relatively high levels of phosphoserine (*23*). Furthermore, recent energy dispersive spectroscopy (EDS) analysis of partially mineralized teeth suggested the presence of phosphorus coexisting with iron oxide (*14*).

The results of immunohistochemical staining showed that the fluorescence signal of AjRTMP was stronger inside the epithelial cells than in the cusps (Fig. 2E and Fig. 3B and C). Like enamel matrix proteins, AjRTMP1 may be compartmentalized in secretory vesicles within epithelial cells (*24*). The iron-free granules reported from previous TEM observations of radular epithelial cells may be secretory vesicles that encapsulate AjRTMP1 (*19*). In addition, the protein band size of AjRTMP1 extracted from the cusp was smaller than that of AjRTMP1 within the epithelial cells (Fig. 2C and D). These findings suggest that after secretion, AjRTMP1 may undergo proteolytic processing similar to that of enamel matrix proteins (*24, 25*).

The amount of F-actin increased, and actin filamentation progressed as the tooth matured (Fig. 3D). Previous studies have reported an increase in the number and length of microvilli with tooth mineralization (*19*). Confocal microscopic analysis confirmed that AjRTMP1 was localized at the tips, bases, and interspaces of F-actin filaments (Fig. 3C). In ameloblasts involved in enamel formation, the F-actin network present at the apical surface has been reported to play a critical role in the transport of vesicles containing enamel matrix proteins (EMPs), such as amelogenin, and the secretion of EMPs. Moreover, a previous report showed that the F-actin network develops on the apical surface during EMP secretion and enamel formation (*26*). Like those in ameloblasts, the F-actin filaments between radular epithelial cells and cusps may serve as pathways or support the movement of secretory vesicles containing AjRTMP1 during transport.

On the basis of the above results, we propose a model in which RTMP1 controls iron oxide deposition on chiton teeth (Fig. 5). First, RTMP1 expressed in radular epithelial cells is secreted extracellularly and binds to chitin fibers at the leading edge of the cusp interior (tooth #−2∼#0). Next, iron ions released from ferritin accumulated around the tooth are deposited as ferrihydrite in the interior of the cusp through interactions with preexisting RTMP1 (tooth #1). Subsequently, RTMP1 is continuously secreted from the cells at the trailing edge of the tooth into the interior of the cusp, where it binds to the chitin just behind the region where iron oxide has been deposited at the leading edge. Iron ions released from ferritin are then deposited, resulting in the layering of iron oxide in the interior of the tooth (tooth #1∼). As the tooth matures, the iron oxide initially deposited on the leading edge crystallizes into magnetite.

**Fig. 5.**
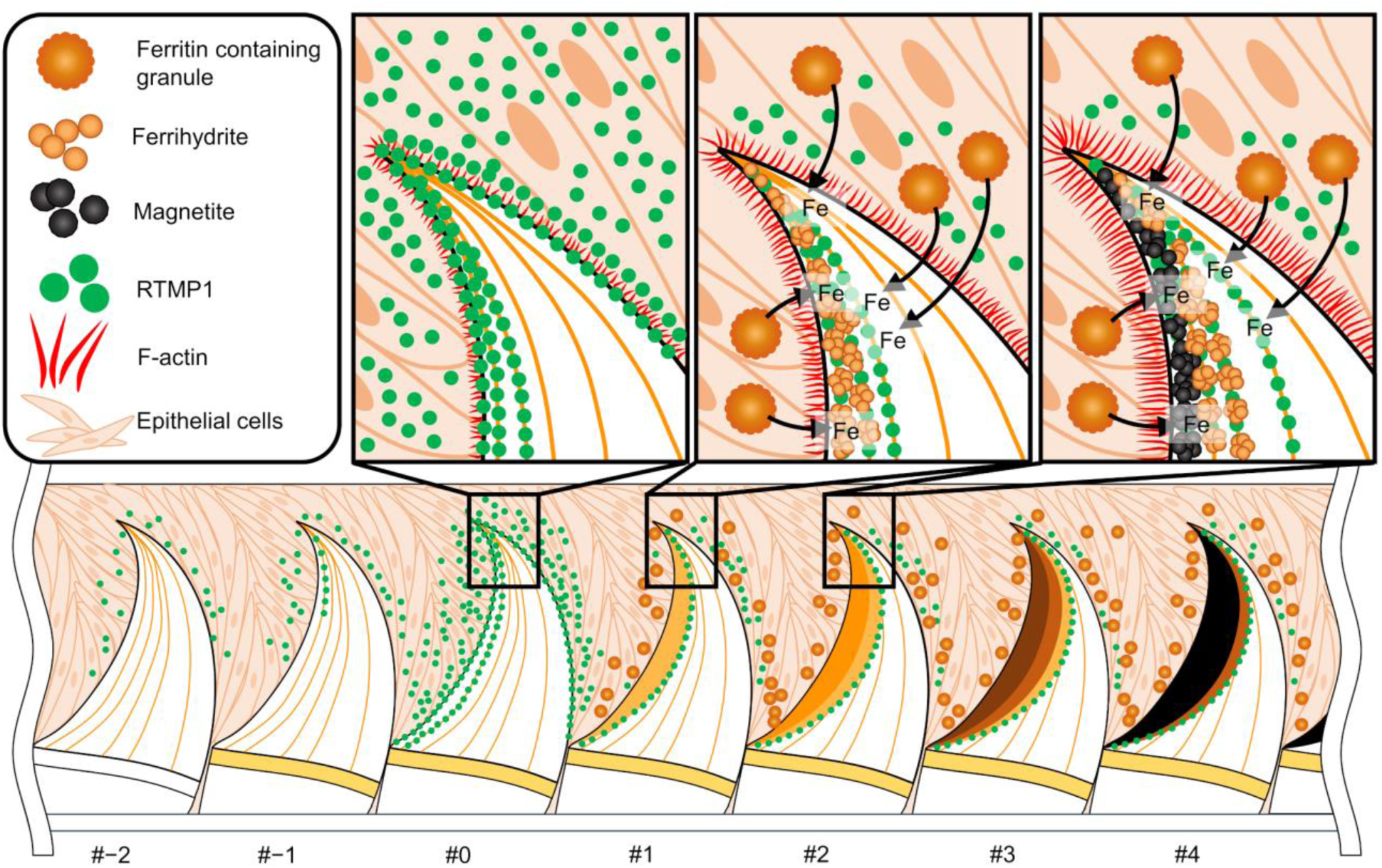
Proposed model of iron oxide deposition on chiton teeth controlled by RTMP1.

## Materials and Methods

### Animals

Live specimens of *Acanthopleura japonica*, *Acanthochitona achates*, and *Placiphorella stimpsoni* were collected from areas in the intertidal zone around Ushimado Harbour, Okayama, Japan, outside protected water areas. Live specimens of *Cryptochiton stelleri* were collected from Akkeshi, Hokkaido, Japan, with special permission from Hokkaido Prefecture, Japan. Animal experiments on live specimens of chitons were conducted in accordance with the guidelines of the Animal Care and Use Committee of Okayama University, and chitons were handled in accordance with procedures approved by the Animal Care and Use Committee of Okayama University (permission no. OKU−2023651).

### RNA isolation and cDNA library construction

Radulae were collected from freshly dissected *A. japonica*, *A. achates*, and *P. stimpsoni*. Radular tissue covered with epithelial cells was extracted from radulae under a stereomicroscope Stemi DV4 (Zeiss, Germany), immediately snap frozen in liquid nitrogen, and stored at −80°C until use. Total RNA was extracted from radular tissues using TRI reagent (Molecular Research Center, Cincinnati, OH, USA) and then purified via the Monarch Total RNA Miniprep Kit (New England Biolabs, Ipswich, MA, USA) following the manufacturer’s instructions. For sufficient RNA, radular tissues isolated from multiple individuals were mixed and subjected to RNA extraction. In this study, two individuals of *A. japonica*, fourteen individuals of *A. achates* and seventeen individuals of *P. stimpsoni* were used for RNA extraction. The integrity and quantity of the extracted RNA were confirmed using the Agilent 2100 Tape station (Agilent Technologies, Santa Clara, CA, USA). The cDNA libraries of the radular tissues were prepared using TruSeq Stranded mRNA Sample Prep Kits (Illumina, San Diego, CA, USA).

### RNA sequencing and de novo transcriptome assembly

The constructed cDNA libraries were sequenced on an Illumina NovaSeq 6000 platform to generate paired-end raw reads. Error correction of the raw reads was performed using Rcorrector (*27*), and erroneous k-mers were discarded using TranscriptomeAssemblyTools (https://github.com/harvardinformatics/TranscriptomeAssemblyTools). The parameters for Rcorrector were set to default values. Adapter sequences and low-quality reads were removed by TrimGalore! (ver. 0.6.5). Low-quality reads with Phred scores of less than 5 were removed. Transcriptome de novo assembly was performed using Trinity (ver. 2.8.6) (*28*). Assembly conditions were performed with default settings, except that normalization was not performed. ORFs were predicted from contigs by TransDecoder (ver. 5.5.0) with the default settings. CD-Hit (*29*) was used to remove highly homologous sequences from ORFs. The Markov clustering threshold for CD-Hit was set at 95%.

### Bioinformatic analysis

Orthofinder was used to estimate the orthologous protein groups between *C. stelleri*, *A. japonica*, *A. achates* and *P. stimpsoni.* A homology search was conducted using BLAST+ (ver. 2.7.1) (*30*). Multiple protein sequence alignment of RTMP1 proteins was performed using T-coffee expresso (*31*). Tandem repeat sequences in RTMP1 were detected using XSTREAM (*32*). The signal peptide was predicted by SignalP (ver.5.0) (*33*). IUPred (*34*) was used to predict intrinsically disordered regions of RTMP1 and its homologs. The TMM-FPKM used to compare the gene expression levels of the mineralized cusp-specific protein was calculated according to the method of Fukushima and Pollock (*35*). Briefly, the cleaned read data used for the de novo transcriptome assembly were mapped to the transcriptome data by Bowtie2 (*36*). Mapping was performed using the options [--dpad 0 --gbar 99999999 --mp 1,1 --np 1 -- score-min L,0,-0.1 --no-mixed --no-discordant]. The resulting SAM file was converted to a BAM file using SAMtools (*37*). Count data were obtained by RSEM (*38*) with the resulting BAM file and the transcriptome data. RSEM was performed with the default settings. The isoforms that mapped to less than 25% of the total were removed. Trimmed mean of M values (TMM) normalization was used to normalize expression levels across species. The scaling factors were determined from the expression levels of the single-copy ortholog groups identified above, and TMM-FPKM was calculated by edgeR (*39*). MAFFT was used for multiple alignment to construct the phylogenetic tree. CO1 sequences of *Chiton olivaceus* (XBY88089.1), *Acanthopleura granulata* (QXE46576.1), *A. japonica* (*Liolophura japonica*) (BDO47095.1), *Ischnochiton australis* (UEF90774.1), *Ischnochiton comptus* (BAV14531.1), *A. achates* (BDO47092.1), *Acanthochitona defilippii* (UGZ24807.1), *C. stelleri* (BDO47091.1), *Placiphorella atlantica* (ADH41916.1) and *P. stimpsoni* (AXG25339.1) were used for the phylogenetic analyses. Multiple sequence alignments were performed using MAFFT (*40*). The alignments were trimmed with TrimAl (ver. 1.4.1) (*41*). RAxML (*42*) was used to construct a phylogenetic tree by maximum likelihood (ML) methods. The best model for phylogenetic analysis was selected by the Protgammaauto option of RaxML. The bootstrap analysis was carried out with 1,000 replications. The phylogenetic tree was drawn with iTOL (ver. 5.0) (https://itol.embl.de/).

### Protein extraction from the radular tissue of *A. japonica*

Protein extraction from the radular tissue of *A. japonica* was conducted in accordance with the methods previously described in our research (*16*). Briefly, thirty-four *A. japonica* individuals were freshly dissected, and their radular tissues were collected. The radular tissue was divided into three stages: stage 1 (8–9 teeth), consisting of transparent teeth composed mainly of chitin; stage 2 (3–5 teeth), consisting of teeth with a reddish-brown color due to iron deposition; and stage 3 (10 teeth), where amorphous iron oxide crystallized into magnetite, forming black teeth. Fractionated radular tissues were combined for each stage and pipetted into 1 mL of PBS, and epithelial cells were separated from the radular tissues. The radulae, comprising teeth, a base, and a radular membrane, were freeze-dried following the removal of epithelial cells. The epithelial cell suspension was centrifuged at 4°C and 13,000 rpm for 1 min to obtain epithelial cell pellets. One hundred microliters of lysis buffer [50 mM Tris (pH 8.0), 1% Triton X- 100, 150 mM NaCl, 2 mM EDTA, 1 mM sodium orthovanadate, 1 mM NaF, 2 µg/mL aprotinin, and 1×EDTA-Free Complete Protease Inhibitor Cocktail (Roche, Mannheim, Germany)] was added to the epithelial cell pellet, which was then sonicated for a period of two minutes on ice. The cell lysate was subjected to centrifugation at 13,000 rpm for one minute at 4°C, after which the supernatant was collected and stored at −20°C until further use. From the freeze-dried radulae, the mineralized cusp, base, and radular membrane were dissected using a scalpel and forceps. The mineralized cusp, base, and radular membrane were rinsed with ultrapure water until no further cellular detritus could be observed. The mineralized cusp, base and radular membranes were washed with 1 mL of 70% ethanol, followed by 1 mL of ultrapure water. The samples were subsequently freeze-dried, weighed, and ground to a fine powder using a mortar and pestle. A 1% sodium dodecyl sulfate (SDS) solution was added to the ground sample, which was subsequently boiled at 100°C for 30 minutes. The mixture was then subjected to centrifugation at 13,000 rpm for one minute at room temperature, after which the supernatant was collected and stored at −20°C until further use. Total protein quantification was performed by a bicinchoninic acid (BCA) assay with a BCA protein assay kit (TaKaRa Bio, Shiga, Japan) following the manufacturer’s instructions.

### Tricine SDS‒polyacrylamide gel electrophoresis

Tricine SDS‒polyacrylamide gel electrophoresis (tricine SDS‒PAGE) was performed according to a previously reported method (*43*). After protein separation, the gels were stained with Coomassie Brilliant Blue R-250.

### Western blotting

The protein samples were subjected to tricine SDS‒PAGE and transferred onto PVDF membranes (Merck Millipore, USA). The membranes were immunoblotted with anti-AjRTMP1 rabbit polyclonal antibody, followed by incubation with goat anti-rabbit IgG H&L (HRP) (Abcam, Cambridge, MA, USA). The anti-AjRTMP1 rabbit polyclonal antibody was generated based on the ORF identified from *A. japonica* transcriptome data. The signals were detected on an ImageQuant LAS 500 (GE Healthcare, Chicago, IL, USA) with Clarity Max™ Western ECL Substrate (Bio-Rad, Hercules, CA, USA).

### Quantitative reverse transcriptase‒polymerase chain reaction (qRT‒PCR) expression analysis

Radular tissue and foot tissue were isolated from freshly dissected *C. stelleri*. After radular tissue isolation, stage 1, stage 2 and stage 3 samples were separated from whole radulae as described above, frozen immediately in liquid nitrogen and stored at −80°C until use. Total RNA was extracted from the radular and foot tissues as described above. The extracted total RNA was converted into first-strand cDNA using the Luna Script RT Super Mix Kit (New England Biolabs). Subsequent qPCR analyses were performed using a an Mx3000P QPCR System (Agilent Technologies) with a Luna Universal qPCR Master Mix Kit (New England Biolabs). The target mRNA expression levels were normalized to those of the actin gene. Relative expression levels were calculated using the ΔΔCt method. All primers used for qRT‒PCR are listed in table S4.

### Paraffin-embedded section preparation of radular tissue from *A. japonica*

Live specimens of *A. japonica* were freshly dissected, and the radular tissue was isolated. The tissue was immediately fixed in 4% paraformaldehyde in PBS for 6 h at 4°C. The sample was then serially dehydrated in ethanol up to 100%, embedded in paraffin, and used to prepare 4-μm-thick tissue sections. The sections were dried using an air dryer.

### Frozen section preparation of radular tissue from *A. japonica*

Live specimens of *A. japonica* were freshly dissected, and the radular tissue was isolated. The tissue was immediately fixed in 4% paraformaldehyde in PBS for 6 h at 4°C. Fixed tissues were then placed in 30% sucrose/PBS and treated overnight at 4°C. After the samples were embedded in O.C.T. compound (Sakura Finetek, Tokyo, Japan), 20-μm-thick frozen sections were prepared using a Leica CM1850 cryostat (Nussloch, Germany). The sections were dried via an air dryer.

### Immunofluorescence staining

Immunofluorescence staining of the sections was primarily performed as previously described (*44*). The anti-AjRTMP1 antibody was used as the primary antibody, and donkey anti-rabbit IgG (H+L) labeled with Alexa Fluor™ 488 (Invitrogen, Carlsbad, CA, USA) was used as the secondary antibody. Counterstaining was performed with Alexa Fluor 594-phalloidin (Invitrogen) to stain F-actin and with Hoechst 33342 solution (Dojindo Molecular Technologies, Kumamoto, Japan) to stain chromosomes. Immunofluorescence-stained paraffin sections were captured using an Olympus BX51 fluorescence microscope (Olympus, Tokyo, Japan), and frozen sections were imaged using an Olympus FV3000 confocal laser scanning microscope (Olympus).

### Fluorescence intensity measurement

The distribution of AjRTMP1 and F-actin in the fluorescence microscope images was determined by measuring the fluorescence intensity using ImageJ (NIH) (*45*). The locations where the fluorescence intensity was measured are indicated by arrows in the figures.

### In vitro mineralization experiments using radula extracted from *A. japonica*

Live specimen of *A. japonica* was freshly dissected, and the radular tissues was isolated. The tissue was pipetted in 1 mL of PBS to remove epithelial cells. As a negative control, the tissue was treated by removing the epithelial cells in the same manner described above, followed by boiling in 1% SDS for 30 minutes to remove native proteins. The treated radula was then placed in 5 mM FeCl_3_ and rotated at 4°C at 25 rpm for 7 days. The radula was removed from the iron solution, washed three times with ultrapure water, and fixed in 2.5% glutaraldehyde at 4°C for 6 h. The samples were then serially dehydrated in ethanol to 100% ethanol and embedded in epoxy resin, and ultrathin sections of approximately 80 nm thickness were prepared using an ultramicrotome. The prepared ultrathin sections were placed on a copper grid and negatively stained with 2% uranyl acetate solution. The teeth of #0, which are speculated to have the highest expression of AjRTMP1 on the basis of the immunofluorescence staining results, were observed by TEM. TEM imaging was performed on a HITACHI H-7650 system (Hitachi, Tokyo, Japan) operating at 80 kV.

### Construction of the expression plasmid

The full-length RTMP1 gene sequence was obtained from the constructed transcriptomic data of *C. stelleri.* For expression of recombinant RTMP1 in budding yeast, the RTMP1 gene sequence without the signal sequence (RTMP1 Δss) was codon-optimized for yeast using OPTIMIZER (*46*) and artificially synthesized. For expression of RTMP1 Δss as a GST fusion protein (GST-RTMP1 Δss), the pEG(KT) vector was used. Both RTMP1 Δss and pEG(KT), which were amplified by PCR using KOD-Plus-Neo DNA polymerase (Toyobo, Osaka, Japan), were introduced into yeast cells, and the plasmid pEG(KT)- RTMP1 Δss was constructed by the homologous recombination activity of yeast cells (*47*). Similarly, the plasmid pEG(KT)-RTMP1 ΔG was constructed to express the GST-RTMP1 ΔG. Additionally, the plasmid pEG(KT)-GFP was constructed to express GST-GFP. The sequence of the constructed plasmids was confirmed by DNA sequencing.

### Expression and purification of recombinant proteins

The budding yeast *Saccharomyces cerevisiae* strain BY4741 was used for the expression of GST- RTMP1 Δss, GST-RTMP1 ΔG and GST-GFP. The transformation of *S. cerevisiae* was performed as described previously (*48*). Yeasts were cultivated in synthetic complete (SC) media without leucine (Leu) and uracil (Ura) (SC-LU). A single colony was grown for 48 h at 30°C at 250 rpm in 50-mL flasks that contained 25 mL of SC-LU medium containing 2% glucose. For large-scale cultivation, instead of using a single colony, 1% of the total culture volume from the preculture was added, followed by incubation for 48 h. At 48 h after the start of culture, the cells were harvested by centrifugation at 6,000 rpm for 10 minutes at room temperature. The medium was removed and replaced with an equal volume of SC-LU medium containing 2% galactose to induce protein expression. At 24 h after the induction of protein expression, the cells were collected by centrifugation at 6,000 rpm for 10 minutes at room temperature. The pellet was washed with a volume of phosphate-buffered saline (PBS) equal to half of the culture medium and then centrifuged again at 6,000 rpm for 10 minutes at room temperature to collect the cells. The cell pellet was resuspended in 800 μL of lysis buffer (1× PBS, 0.05% Tween 20, 1 mM DTT, and 1× EDTA-Free Complete Protease Inhibitor Cocktail [Roche]) per 25 mL of culture medium. The cells were disrupted with glass beads using a Precellys24 cell disruption system (Bertin Technologies, Montigny-le-Bretonneux, France) at 5,000 rpm for 30 seconds, which was repeated seven times. The cell extracts were clarified by centrifugation at 15,000 rpm for 5 minutes at 4°C, and the supernatant was collected. The GST fusion proteins were affinity purified using Glutathione Sepharose™ 4B (GE Healthcare) following the batch purification protocol according to the manufacturer’s instructions. The purified protein mixture was dialyzed overnight at 4°C in PBS using a cellulose dialysis tube (Sekisui Medical, Tokyo, Japan). The dialyzed solution was concentrated using an Amicon Ultra-0.5 Centrifugal Filter Unit (3 kDa cutoff) (Merck Millipore). The purity of the purified protein was confirmed by SDS‒PAGE (*49*). The purified protein solutions were stored at 4°C and used in experiments within one week.

### Chitin binding assay

Purified GST-RTMP1 ΔG, GST-GFP, and the commercially obtained Wheat Germ Agglutinin (WGA) (accession no. (Gene/Protein) P10968) (positive control) (Mitsubishi Gas Chemical, Tokyo, Japan) were dialyzed overnight at 4°C in binding buffer (0.5 M NaCl, 10 mM Tris-HCl, 0.05% Triton X-100, pH 7.0) using a cellulose dialysis tube (MWCO 3500). A 200 μL aliquot of protein mixture, adjusted to 0.9 μg/μL with binding buffer, was mixed with 7.4 μL of colloidal chitin solution and adjusted to 30 μg/μL with binding buffer. The mixture was rotated at 4°C and centrifuged at 12,000 rpm for 5 minutes at 4°C, 1 h after mixing, and 10 μL of the supernatant was collected to create the unbound (1 h) fraction. The solution was then resuspended and rotated again at 4°C, and the unbound (3 h) fraction was collected in the same manner 3 h after mixing the colloidal chitin and protein solutions. The unbound (20 h) fraction was further collected in the same way 20 h after mixing. The colloidal chitin pellets obtained 20 h after mixing were washed three times with 300 µL of binding buffer and centrifuged at 12,000 rpm for 5 minutes at 4°C to remove the supernatant. To the colloidal chitin pellet, 20 µL of SDS sample buffer was added to form the bound fraction. Each collected fraction was subjected to SDS‒PAGE (*49*). After protein separation, the gels were stained with Coomassie Brilliant Blue R 250, and the chitin-binding capacity of the proteins was evaluated.

### Iron binding assay

Purified GST-RTMP1 Δss, GST-RTMP1 ΔG, GST-GFP, and commercially obtained horse spleen ferritin (positive control) (Cat # F4503, Sigma‒Aldrich, St. Louis, MO, USA) were dot blotted onto a PVDF membrane. The protein-blotted PVDF membrane was incubated in 5 mM FeCl_3_ solution for 30 minutes at room temperature. Unbound iron was removed by rinsing the membrane three times in PBS at room temperature for 30 minutes. For detection of iron bound to the PVDF membrane, the membrane was stained with a freshly prepared 1:1 (vol/vol) mixture of 2% K_4_Fe (CN)_6_ (ferrocyanide or Prussian blue; Sigma) and 2% 11.6 M HCl (resulting in a final HCl concentration of 0.116 M) for 10 minutes at room temperature. For confirmation of protein fixation on the PVDF membrane, the membrane was blotted in the same manner and stained with Coomassie Brilliant Blue R 250.

### In vitro iron oxide formation on chitin fibers using recombinant RTMP1

Marine nanofiber (CN-Exp19064, Lot No: F219100101), commercial chitin fiber dispersed in water at approximately 2% concentration, was diluted 10-fold with PBS. Twenty microliters each of GST- RTMP1 Δss and GST-GFP, adjusted to 3 μg/μL, were added to 10 μL of the diluted marine nanofiber and mixed. The mixture was rotated at 4°C at 25 rpm for 24 h. The marine nanofibers were collected by centrifugation at 12,000 rpm for 5 minutes at 4°C, and the supernatant was discarded to remove unbound proteins. The marine nanofiber pellet was washed three times with 500 μL of PBS and centrifuged again at 12,000 rpm for 5 minutes at 4°C, after which the marine nanofibers were collected. Five millimolar FeCl_3_ was added to the collected marine nanofibers, and the mixture was rotated at 4°C at 25 rpm for 7 days. For removal of unbound iron from the reaction mixture, the marine nanofibers were washed three times with 500 μL of ultrapure water using an Amicon Ultra-0.5 Centrifugal Filter Unit (3 kDa cutoff) (Merck Millipore). The collected marine nanofibers were dropped onto a copper grid, negatively stained with 2% uranyl acetate solution, and analyzed by STEM-EDX.

### STEM-EDX

Before analysis, the samples were carbon coated with a VC-100S carbon coater (Vacuum Device, Ibaraki, Japan). STEM-EDX analysis was performed using a field-emission scanning electron microscope (FE-SEM) SU9000 (Hitachi) equipped with an X-ray energy dispersive spectroscopy (EDS) system operating at 25 kV.

### Phosphoproteomic analysis

Proteins were extracted from the radular tissue of 10 *C. stelleri* specimens using the same method as described above. The samples extracted from the epithelial cells of stage 2 were subjected to phosphoproteomic analysis. The extracted proteins were separated using tricine-SDS‒PAGE and stained with Coomassie Brilliant Blue R 250. A band near 56 kDa was excised from the gel with a scalpel, followed by in-gel digestion with trypsin, and the peptides were extracted. Phosphopeptides were enriched from the extracted peptides using the HighSelect™ TiO₂ Phosphopeptide Enrichment Kit (ThermoFisher, Waltham, MA, USA) according to the manufacturer’s instructions. The enriched phosphopeptides were analyzed by high-performance liquid chromatography (HPLC)-Chip and quadrupole time‒of‒flight mass spectrometry (QTOF-MS; G6520 and G4240, Agilent Technologies) as previously reported (*50*). The obtained tandem mass spectra (MS/MS) data were searched for phosphorylated peptides using the MASCOT algorithm against the ORF database of *C. stelleri* with the parameters described previously (*15, 16*). Purified GST-RTMP1 Δss proteins were separated by SDS‒ PAGE, and CBB-stained bands were excised and in-gel digested, followed by HPLC‒Chip-QTOF‒MS analysis as described above.

### RNAi experiment

First, the AjRTMP1 gene was cloned using the following method. Total RNA was extracted from the radular tissue of *A. japonica* using the method described above, and cDNA was synthesized using the SuperScript III First-Strand Synthesis System for qRT‒PCR (Invitrogen) according to the manufacturer’s instructions. The synthesized cDNA was then used as a template for PCR amplification of AjRTMP1 cDNA using the primers AjRTMP1dsRNA_F and AjRTMP1dsRNA_R listed in table S5 and TaKaRa Ex Taq (TaKaRa Bio). The resulting PCR product was subsequently cloned and inserted into the pMD20-T vector (TaKaRa Bio) using the Mighty TA-cloning Kit (TaKaRa Bio) following the manufacturer’s instructions. Next, the cloned AjRTMP1 gene was used as a template for PCR, along with the primers AjRTMP1dsRNA_TF and AjRTMP1dsRNA_R or AjRTMP1dsRNA_TR and AjRTMP1dsRNA_F listed in table S5, to generate PCR products with T7 promoter sequences. With these PCR products, AjRTMP1 ssRNA was synthesized using an in vitro transcription T7 kit for siRNA synthesis (TaKaRa Bio) following the manufacturer’s instructions. AjRTMP1 dsRNA was obtained by mixing and annealing the synthesized AjRTMP1 ssRNA. AjRTMP1 dsRNA was treated with DNase I, purified, dissolved in PBS, and stored at −80°C until use. In a similar manner, GFP dsRNA was prepared as a negative control using pLenti CMV GFP Zeo (Addgene plasmid #17,449) as the template, and the primers GFPdsRNA_TF and GFPdsRNA_R, GFPdsRNA_TR and GFPdsRNA_F are listed in table S5.

Live specimens of *A. japonica* were kept in natural seawater for one day. Three groups were set up for injections: 30 μg of AjRTMP1 dsRNA in 50 μL of PBS, 30 μg of GFP dsRNA in 50 μL of PBS, and 50 μL of PBS. Five individual chitons were used for each group. The dsRNA was injected through the gills using a 1 mL syringe with a 27G needle (Terumo, Tokyo, Japan), taking care not to damage the internal organs. After the dsRNA was injected, the chitons were kept in natural seawater for 48 h and then dissected, after which the radular tissues were collected. One of the two rows of radular tissue was used for gene expression analysis, and the other row was used for phenotypic analysis. Total RNA was extracted as described above, and qRT‒PCR was performed using the primers listed in table S5 (AjRTMP1_real_F and AjRTMP1_real_R, Ajreal_Actin_F and Ajreal_Actin_R) and the method described above to measure the relative expression levels of AjRTMP1. For phenotypic analysis, paraffin sections were prepared as described above and observed using a BZ-X710 microscope (KEYENCE, Osaka, Japan). A square area (10×10 μm) in the center of the cusps of each sample was enclosed, and the mean saturation value was measured. The relative values were calculated by setting the saturation value of tooth #1 to 1 for each sample, and these relative values were plotted.

### Statistical analysis

The Kruskal‒Wallis test was used to compare gene expression levels among the three groups in the RNAi experiments. For comparisons between individual groups, the Mann‒Whitney U test was employed with correction using the Bonferroni method. A significance level of 5% (p < 0.05) was used to determine statistical significance for all tests.

### Availability of data and materials

The reads derived from the Illumina NovaSeq 6000 platform for the analyzed samples have been deposited in the DDBJ Sequence Read Archive under accession numbers, DRR598739, DRR598778 and DRR598779.

## Acknowledgments

We thank S. Hamano of the Akkeshi Marine Station, Field Science Center for Northern Biosphere, Hokkaido University, and W. Godo and K. Saito of the Ushimado Marine Institute, Okayama University, for their assistance in collecting live specimens of chitons. We also thank M. Owada of the Faculty of Science, Department of Biological Sciences, Kanagawa University, for helpful advice regarding the identification of the chiton species used in this study. We are grateful to M. Monobe and T. Tsukano at the Central Research Laboratory of Okayama University for their technical support in preparing paraffin-embedded and epoxy resin-embedded ultrathin sections. We thank T. Shiokawa and H. Tada at the Division of Instrumental Analysis for their assistance with the LC‒ MS/MS measurements and I. Nakamura of Okayama University for help with sample preparation for these measurements. Special thanks to Chiyu Nakano at the Organization for Research Strategy and Development, Okayama University, for technical support with STEM-EDX analysis. We also thank S. Ifuku of Kyoto University for providing marine nanofibers and T. Nitoda of Okayama University for providing colloidal chitin. We appreciate the insightful discussions with K. Aoki of Okayama University of Science, H. Yamada of Okayama University, J. A. Shaw of the University of Western Australia, and M. Suzuki of the University of Tokyo. Computations were partially performed on the NIG supercomputer at the ROIS National Institute of Genetics. We also thank American Journal Experts (AJE) for their assistance with language editing.

## Funding

Ministry of Education, Culture, Sports, Science, and Technology (MEXT Japan) KAKENHI 21K05781 (MN)

Ministry of Education, Culture, Sports, Science, and Technology (MEXT Japan) KAKENHI 22H04812 (MN)

Japan Science and Technology Agency (JST) Fusion Oriented REsearch for disruptive Science and Technology (FOREST) JPMJFR210E (MN)

JGC Saneyoshi Scholarship Foundation (MN) Kato Memorial Bioscience Foundation (MN)

## Author contributions

Conceptualization: M N, K Okada

Funding acquisition: M N

Investigation: M N, K Okada, H A, Y O, Y N, K Okoshi, K Obuse, D K, H M and A S

Software: K Obuse

Supervision: M N

Writing – original draft: M N, K Okada

Writing – review & editing: M N, K Okada, K Okoshi, K Obuse, D K, H M and A S

## Data and materials availability

RNA-seq data have been deposited in the DDBJ Sequence Read Archive under accession numbers, DRR598739, DRR598778 and DRR598779.

