## Supplementary Materials for "Radular teeth matrix protein 1 directs iron oxide deposition in chiton teeth"

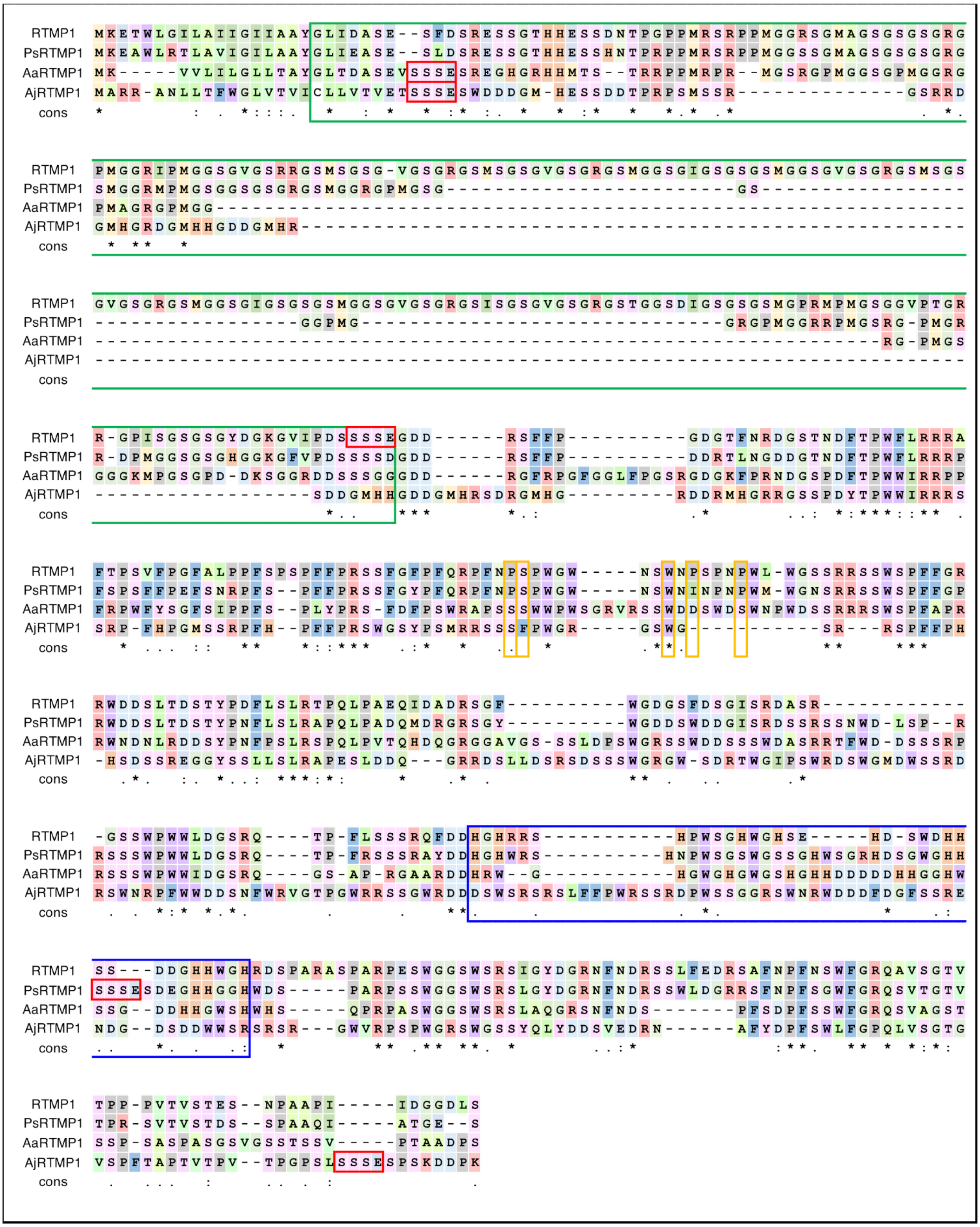
RTMP1 MKETWLGILAIIGIIAAYGLIDASE--SFDSRESSGTHHESSDNTPGPPMRSRPPMGGRSGMAGSGSGSGRG
PsRTMP1 MKEAWLRTLAVIGILAAYGLIEASE--SLDSRESSGTHHESSHNTPRPPMRSRPPMGGSSGMAGSGSGSGRG
AaRTMP1 MK-----VVLILGLLTAYGLTDASEVSSSESREGHGRHHMTS--TRRPPMRPR--MGSRGPMGGSGPMGGRG
AjRTMP1 MARR-ANLLTFWGLVTVICLLVTVETSSSESWDDDGM-HESSDDTPRPSMSSR-------------GSRRD
cons
RTMP1 PMGGRIPMGGSGVGSRRGSMSGSG-VGSGRGSMSGSGVGSGRGSMGGSGIGSGSGSMGGSGVGSGRGSMSGS
PsRTMP1 SMGGRMPMGSGGSGSGRGSMGGRGPMGSG-----------------------GS------------------
AaRTMP1 PMAGRGPMGG--------------------------------------------------------------
AjRTMP1 GMHGRDGMHHGDDGMHR-------------------------------------------------------
cons
RTMP1 GVGSGRGSMGGSGIGSGSGSMGGSGVGSGRGSISGSGVGSGRGSTGGSDIGSGSGSMGPRMPMGSGGVPTGR
PsRTMP1 ----------------GGPMG-----------------------------GRGPMGGRRPMGSRG-PMGR
AaRTMP1 ---------------------------------------------------------------RG-PMGS
AjRTMP1 ------------------------------------------------------------------------
cons
RTMP1 R-GPISGSGSGYDGKGVIPDSSSSEGDD------RSFFP----------GDGTFNRDGSTNDFTPWFLRRRA
PsRTMP1 R-DPMGGSGSGHGGKGFVPDSSSSDGDD------RSFFP----------DDRTLNGDDGTNDFTPWFLRRRP
AaRTMP1 GGGKMPGSGPD-DKSGGRDDSSSGGGDD------RGFRPGFGGLFPGSRGDGKFPRNDGSPDFTPWWIRRPP
AjRTMP1 -----------------SDDGMHHGDDGMHRSDRGMHG--------RDDRMHGRRGSSPDYTPWWIRRRS
cons
RTMP1 FTPSVFPGFALPPFSPSPFFPRSSFGFPFQRPFNPSPWGW-----NSWNPSPNPWL-WGSSRRSSWSPFFGR
PsRTMP1 FSPSFFPEFSNRPFS--PFFPRSSFGYPFQRPFNPSPWGW-----NSWNINPNPWM-WGNSRRSSWSPFFGP
AaRTMP1 FRPWFYSGFSIPPFS--PLYPRS-FDFPSWRAPSSSWWPWSGRVRSSWDDSWDSWNPWDSSRRRSWSPFAPR
AjRTMP1 SRP-FHPGMSSRPFH--PFFPRSWGSYPSMRRSSSFPWGR-----GSWG-----------SR---RSPFFPH
cons
RTMP1 RWDDSLTDSTYPDFLSLRTPQLPAEQIDADRSGF---------WGDGSFDSGISRDASR-----------
PsRTMP1 RWDDSLTDSTYPNFLSLRAPQLPADQMDRGRSGY---------WGDDSWDDGISRDSSRSSNWD-LSP--R
AaRTMP1 RWNDNLRDDSYPNFPSLRSPQLPVTQHDQGRGGAVGS-SSLDPSWGRSSWDDSSSWDASRRTFWD-DSSSRP
AjRTMP1 -HSDSSREGGYSSLLSLRAPESLDDQ---GRRDSLLDSRSDSSSWGRGW-SDRTWGIPSWRDSWGMDWSSRD
cons
RTMP1 -GSSWPWWLDGSRQ----TP-FLSSSRQFDDHGHRRS-----------HPWSGHWGHSE-----HD-SWDHH
PsRTMP1 RSSSWPWWLDGSRQ----TP-FRSSSRAYDDHGHWRS----------HNPWSGSWGSSGHWSGRHDSGWGHH
AaRTMP1 RSSSWPWWIDGSRQ----GS-AP-RGAARDDHRW--G-----------HGWGHGWGSHGHHDDDDDHHGGHW
AjRTMP1 RSWNRPFWWDDSNFWRVGTPGWRRSSGWRDDDSWSRSRSLFFPWRSSRDPWSSGGRSWNRWDDDFDGFSSRE
cons
RTMP1 SS---DDGHHWGHRDSPARASPARPESWGGSWSRSIGYDGRNFNDRSSLFEDRSAFNPFNSWFGRQAVSGTV
PsRTMP1 SSSESDEGHHGGHWDS-----PARPSSWGGSWSRSLGYDGRNFNDRSSWLDGRRSFNPFSGWFGRQSVTGTV
AaRTMP1 SSG--DDHHGWSHWHS-----QPRPASWGGSWSRSLAQGRSNFNDS-------PFSDPFSSWFGRQSVAGST
AjRTMP1 NDG--DSDDWWSRSRSR---GWVRPSPWGRSWGSSYQLYDDSVEDRN------AFYDPFSWLFGPQLVSGTG
cons
RTMP1 TPP-PVTVSTES--NPAAPI-----IDGGDLS
PsRTMP1 TPR-SVTVSTDS--SPAAQI-----ATGE--S
AaRTMP1 SSP-SASPASGSVGSSTSSV-----PTAADPS
AjRTMP1 VSPFTAPTVTPV--TPGPSLSSSESPSKDDPK
cons

**Fig. S1.**

**Amino acid sequence alignment of RTMP1 and its homologs.** Putative chitin-binding residues are indicated by orange boxes. SSSE motif are indicated by red boxes. G-rich repeats are indicated by green boxes. H-rich domains are indicated by blue boxes.

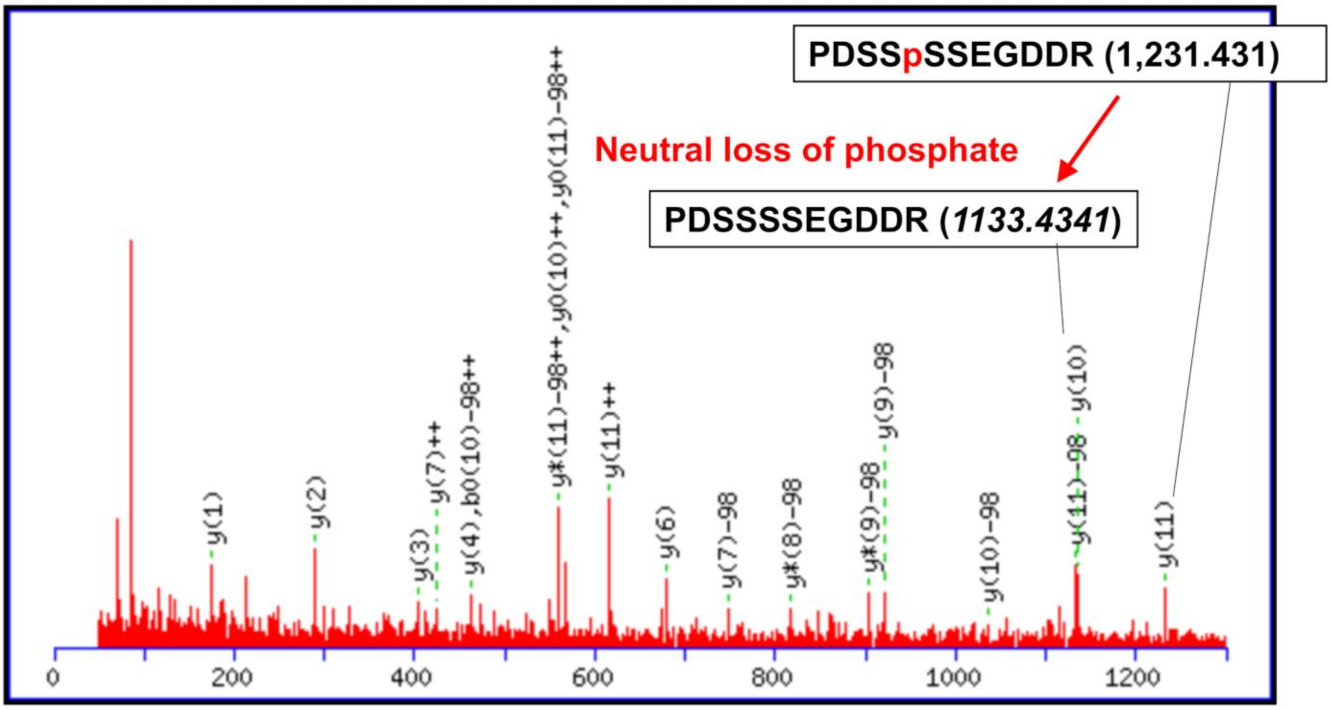


**Fig. S2.**

**Fragmentation spectra of the phosphopeptide of sequence PDSSpSSEGDDR in RTMP1.**

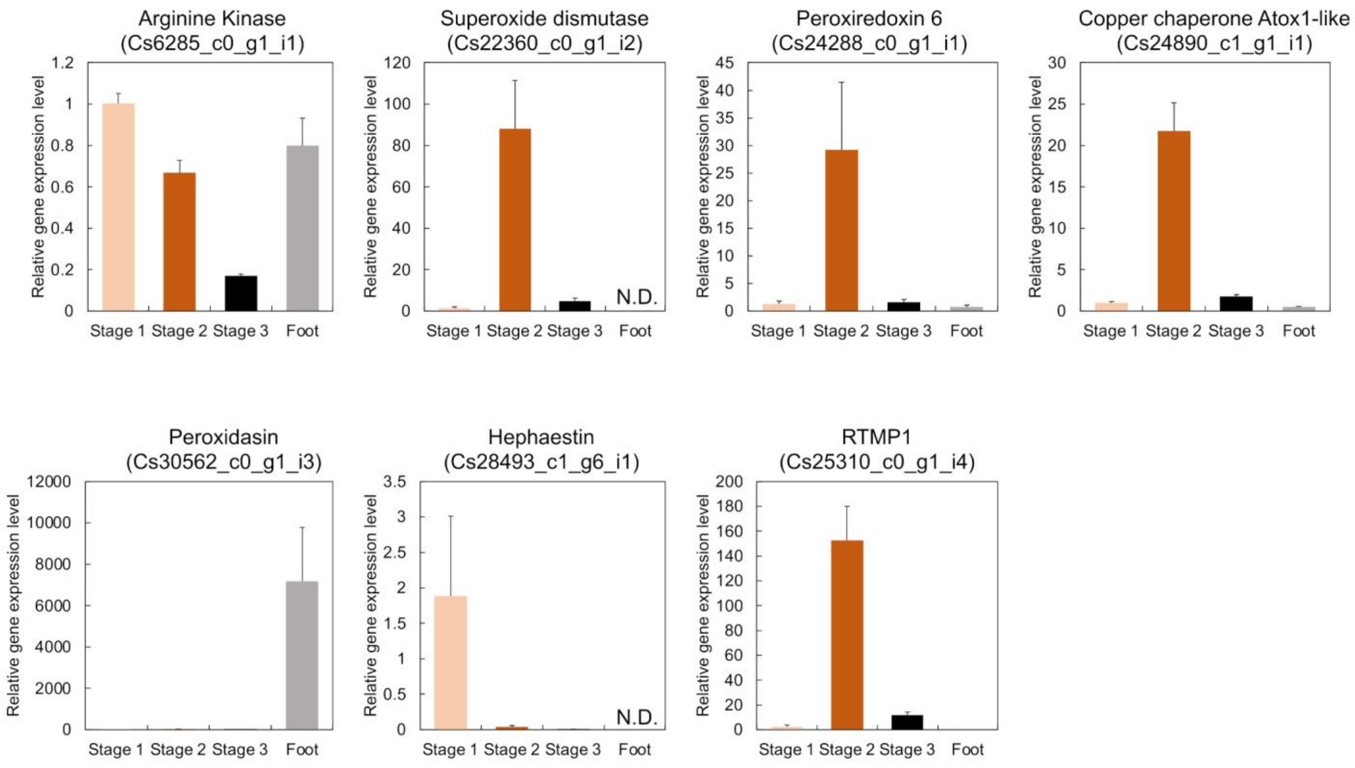


**Fig. S3.**

**qRT–PCR was performed to determine the gene expression profiles of the mineralized cusp-specific proteins during teeth formation in the radular tissue of *C. stelleri*.** The radular tissue was divided into three stages according to the degree of mineralization (Fig. 2A and B). Total RNA was extracted from each stage of radular tissue. Foot tissue was used as a control. Peroxiredoxin 6, superoxide dismutase, copper chaperone Atox1 and RTMP1 were most highly expressed in stage 2. Hephaestin was most highly expressed in stage 1. Among the three peroxidasins identified as mineralized cusp-specific proteins, the expression profile of Cs30562_c0_g1_i3, which presented the highest TPM expression level, was analyzed. The expression level of this gene was much greater in foot tissue than in radular tissue. The expression levels of arginine kinases in the radular and foot tissues were comparable. The data represent the means + standard errors of the means (SEMs) of triplicate biological samples from three independent experiments.

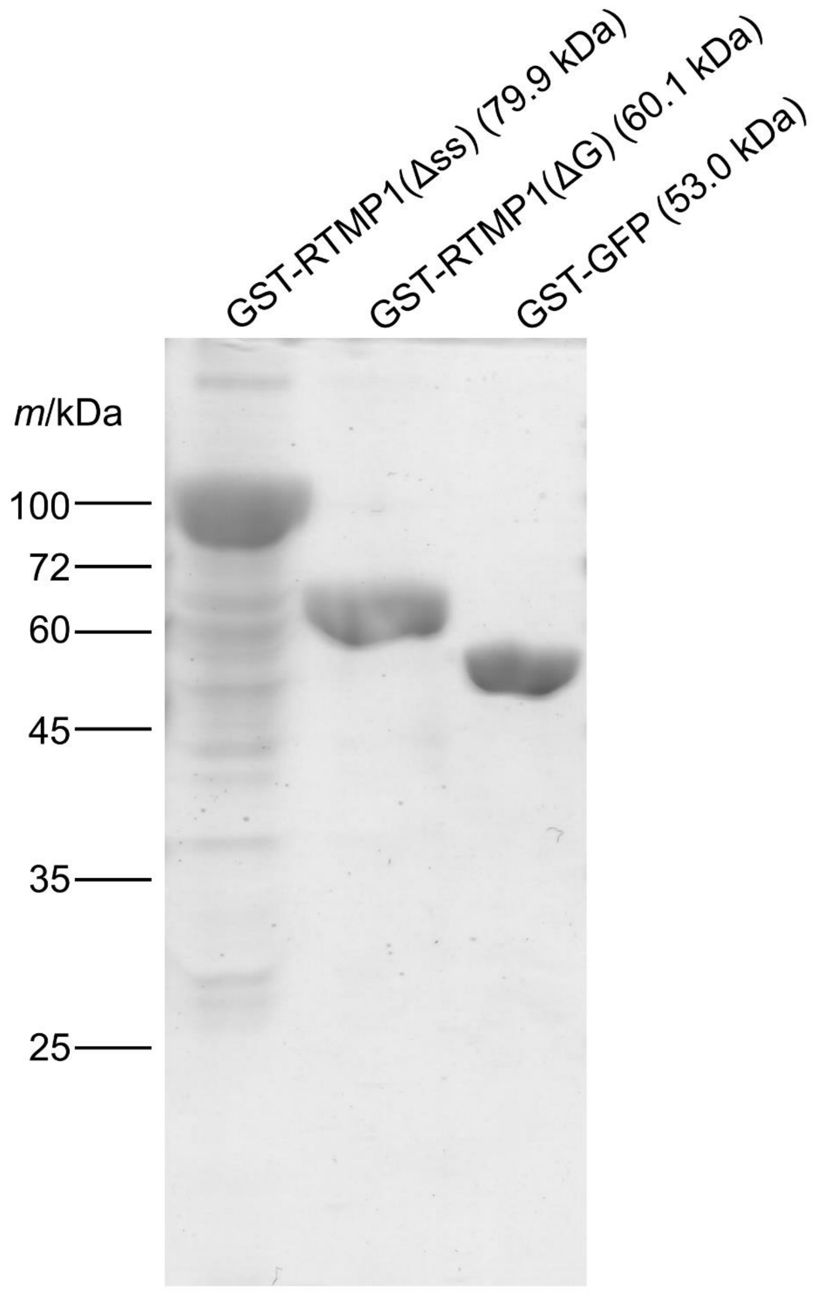


**Fig. S4.**

**SDS–PAGE analysis of recombinant GST-RTMP1 Δss, GST-RTMP1 ΔG and GST-GFP.**

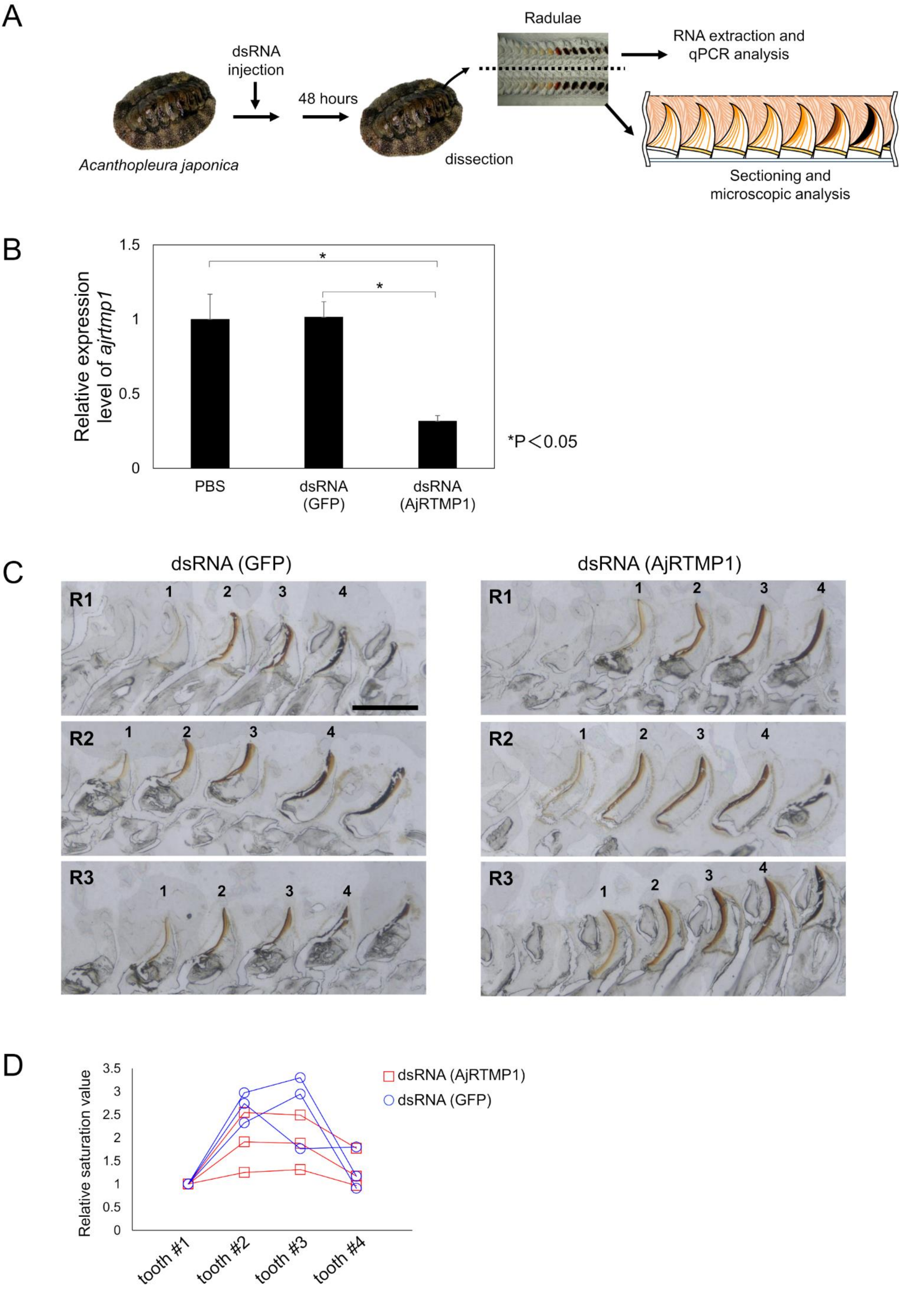
A
dsRNA injection
48 hours
Acanthopleura japonica
dissection
Radulae
RNA extraction and qPCR analysis
Sectioning and microscopic analysis
B
Relative expression level of ajrtmp1
1.5
1
0.5
0
*
*
*P＜0.05
PBS
dsRNA (GFP)
dsRNA (AjRTMP1)
C
dsRNA (GFP)
dsRNA (AjRTMP1)
R1
R2
R3
1
2
3
4
D
Relative saturation value
3.5
3
2.5
2
1.5
1
0.5
0
tooth #1
tooth #2
tooth #3
tooth #4
dsRNA (AjRTMP1)
dsRNA (GFP)

**Fig. S5.**

**Results of the RNAi experiment using *A. japonica*.** (**A**) Schematic of the RNAi experiment. (**B**) The expression levels of AjRTMP1 mRNA in the radular tissue were determined with real-time quantitative PCR 48 h after injection. The AjRTMP1 mRNA expression level in the PBS group was assigned a relative value of 1.0. Five chitons (n = 5) were used in each experiment. Statistically significant differences were identified using the Kruskal–Wallis test. For comparisons between individual groups, the Mann–Whitney U test was used, with the Bonferroni correction applied. (**C**) A bright field microscopic image of the radular tissue from the chitons injected with GFP dsRNA and AjRTMP1 dsRNA. Scale bars, 300 µm. (**D**) The saturation value of the color of the cusps in the microscopic images. A square area (10×10 µm) in the center of the cusps of each sample was enclosed, and the mean saturation value was measured. The relative values were calculated by setting the saturation value of tooth #1 to 1 for each sample, and these relative values were plotted. Iron deposition initially results in the formation of cusps with a low-saturation light brown color. As iron deposition progresses and the brown color darkens, the saturation increases. As the iron deposition continues and matures, the cusps become a black color with low saturation.

**Table S1.**

Summary of RNA-seq, de novo assembly and ORF predictions of chitons.

| Chiton name | *C. stelleri* | *A. japonica* | *A. achates* | *P. stimpsoni* |
|---|---|---|---|---|
| Number of raw reads (PE) | 88,943,700 | 85,061,132 | 79,132,504 | 67,216,174 |
| Number of contigs | 168,867 | 145,822 | 118,026 | 107,613 |
| N50 | 1,374 | 1,892 | 1,546 | 1,012 |
| Number of ORFs | 54,236 | 46,069 | 38,785 | 40,745 |

**Table S2.**

Summary of homologous sequence searches of 22 mineralized cusp-specific proteins identified in *C. stelleri*.

| *C. stelleri* | | | *A. japonica* | | *A. achates* | | *P. stimpsoni* | |
|---|---|---|---|---|---|---|---|---|
| Transcript ID * | annotation | TMM-FPKM | E-value | TMM-FPKM | E-value | TMM-FPKM | E-value | TMM-FPKM |
| Cs30238_c2_g1_i1.p1 | Neuroglobin-like | 651.6 | 8E-64 | 3.5 | 2E-69 | 3.9 | 7E-76 | 1121.5 |
| Cs25764_c0_g1_i1.p1 | Myoglobin | 980.4 | 8E-93 | 4163.2 | 6E-84 | 14886 | 7E-98 | 522 |
| Cs14014_c0_g1_i1.p1 | Protein usf-like | 282.8 | 1E-151 | 169.2 | 8E-124 | 24.7 | 7E-150 | 866.6 |
| Cs6285_c0_g1_i1.p1 | Arginine kinase | 345.9 | 0 | 74.8 | 0 | 1012.3 | 0 | 794.4 |
| Cs30562_c0_g1_i3.p1 | Peroxidasin | 5.1 | 4E-120 | 2.9 | 0 | 3.7 | 0 | 3.4 |
| Cs25310_c0_g1_i4.p1 | RTMP1 (Radular Teeth Matrix Protein 1) | 37.6 | 0.000001 | 218.6 | 2E-22 | 303.5 | 9E-118 | 332.8 |
| Cs7512_c0_g1_i1.p1 | Protein yellow-like | 9.3 | 1E-109 | 11.9 | 0 | 5.8 | 0 | 2.9 |
| Cs30562_c0_g1_i13.p1 | Peroxidasin | 1.3 | 4E-13 | 5.3 | 2E-29 | 1.2 | 4E-60 | 3.4 |
| Cs29973_c1_g11_i2.p1 | Gastric intrinsic factor-like protein 2 | 11.9 | 1E-49 | 0.6 | 7E-46 | 57.1 | 3E-40 | 0.5 |
| Cs28145_c2_g1_i3.p1 | Apoptosis inducing factor-3 | 25.8 | 0 | 104.8 | 0 | 72.8 | 0 | 4.9 |
| Cs24890_c1_g1_i1.p2 | Copper chaperone Atox1-like | 619.7 | 1E-33 | 372.2 | 1E-24 | 529 | 3E-40 | 1246.1 |
| Cs27589_c0_g2_i1.p1 | Hillarin | 14 | 0 | 26.8 | 0 | 41.6 | 0 | 57.7 |
| Cs27160_c6_g1_i10.p1 | Muscle-specific protein 20-like isoform X2 | 52.1 | 2E-117 | 45.1 | 4E-119 | 85.9 | 2E-125 | 87.5 |
| Cs16206_c0_g1_i1.p1 | Eukaryotic initiation factor 4A-II, partial | 68 | 0 | 32.1 | 0 | 146.2 | 0 | 132.6 |
| Cs30504_c2_g1_i3.p1 | Histone H2B 8-like | 2.4 | 4E-80 | 1.9 | 4E-80 | 2.3 | 1E-86 | 1.9 |
| Cs28493_c1_g6_i1.p1 | Hephaestin-like protein | 23.1 | 0 | 43.8 | 0 | 53.8 | 0 | 474.7 |
| Cs29020_c0_g2_i1.p1 | Filamin-A-like isoform X1 | 30.6 | 0 | 20.9 | 0 | 19.3 | 0 | 17.9 |
| Cs24288_c0_g1_i1.p1 | Peroxiredoxin 6 | 132.9 | 5E-137 | 107.2 | 7E-133 | 148 | 2E-143 | 98.2 |
| Cs29504_c3_g1_i2.p1 | Peroxidasin | 2.1 | 1E-112 | 2.9 | 0 | 7.7 | 0 | 3.4 |
| Cs22360_c0_g1_i2.p1 | Superoxide dismutase (Cu-Zn) | 14.5 | 5E-106 | 34.3 | 4E-129 | 118.4 | 5E-144 | 100.1 |
| Cs25988_c1_g3_i1.p1 | Filamin-A-like isoform X5 | 4.4 | 0 | 20.9 | 0 | 14.8 | 0 | 17.9 |
| Cs29440_c6_g2_i1.p1 | Cytosolic malate dehydrogenase | 26.9 | 0 | 23.9 | 0 | 133.8 | 0 | 56.2 |

*Newly assigned TranscriptID in this study.

**Table S3.**

Phosphorylated peptides identified from recombinant GST-RTMP1 Δss.

| Mascot scores | Peptides |
|---|---|
| 59 | GpSMSGSGVGSGR |
| 74 | GSpTGGSDIGSGSGSMGPR |
| 68 | GpSTGGpSDIGSGSGSMGPR |
| 80 | GSMGGpSGIGSGSGSMGGSGVGSGR |
| 75 | GpSMGGSGIGSGSGSMGGSGVGSGR |

**Table S4.**

Sequences of the primers used for qRT–PCR in this study.

| Target gene | Primer Sequence (5' to 3') |
|---|---|
| Cs28493_c1_g6_i1 | FWD: TCTTCGGTAACACCCTGTCC |
| | REV: TCCACCGCACCAACATAGTA |
| Cs6285_c0_g1_i1 | FWD: GCCCGATTACCGAAGTTGGG |
| | REV: CCAGACGGCGTTTGTTGGAG |
| Cs30562_c0_g1_i3 | FWD: GCCCCAGTAGCAGATTTGGC |
| | REV: GGTTGTCCTCAGCAGACCTC |
| Cs24890_c1_g1_i1 | FWD: TGTCTGCACAGAAGCACGAG |
| | REV: AGCAACACCCACATAACTGG |
| Cs24288_c0_g1_i1 | FWD: TAGCACTCACAGCCAGAGC |
| | REV: CTTGACGTCCACGCCCTTAG |
| Cs22360_c0_g1_i2 | FWD: CACGGGAGGATGTGCATCTG |
| | REV: CCCAACCCAAGGTCGTCTTC |
| Cs25310_c0_g1_i4 | FWD: AGCTGGTCCAGAAGCATAGG |
| | REV: GACTGTTCCCGACACAGCTT |
| Actin gene | FWD: CCAGGTATCGCTGACCGTAT |
| | REV: CACCGATCCAGACGGAGTAT |

**Table S5.**

Sequences of the primers used for the RNAi experiment in this study

| Primer name | Primer Sequence (5' to 3') |
|---|---|
| AjRTMP1dsRNA_TF | FWD: GCGTAATACGACTCACTATAGGGAGAAACCTGTTGACGTTCTGGG |
| AjRTMP1dsRNA_R | REV: GAGCTGTCGGAGTGATGAG |
| AjRTMP1dsRNA_TR | REV: GCGTAATACGACTCACTATAGGGAGAGAGCTGTCGGAGTGATGAG |
| AjRTMP1dsRNA_F | FWD: AACCTGTTGACGTTCTGGG |
| GFPdsRNA_TF | FWD: GCGTAATACGACTCACTATAGGGAGAGGGCGAGGAGCTGTTCAC |
| GFPdsRNA_R | REV: CAGGTAGTGGTTGTCGGGC |
| GFPdsRNA_TR | REV: GCGTAATACGACTCACTATAGGGAGACAGGTAGTGGTTGTCGGGC |
| GFPdsRNA_F | FWD: GGGCGAGGAGCTGTTCAC |
| AjRTMP1_real_F | FWD: CCTACCAGCTTTACGACGACA |
| AjRTMP1_real_R | REV: TCCTTAGAGGGCGACTCTGAG |
| Ajreal_Actin_F | FWD: AACCACAGCCGAGAGAGAAA |
| Ajreal_Actin_R | REV: ACCTGAATCTCTCGTTGCCA |
